# P2X7 receptor antagonism potentiates seizure-suppressive effects of anti-seizure medications in human resected epileptic brain tissue

**DOI:** 10.64898/2026.09.18.752730

**Authors:** Ana Fernandez Martin, Beatriz Gil, Meghma Mitra, Jaideep Kesavan, Austin Lacey, Meng-Juan Sun, Anne Marie O’Callaghan, Vincent Healy, Norman Delanty, Alan Beausang, Francesca M. Brett, Michael A. Farrell, Jane Cryan, Donncha F. O’Brien, Kieron J. Sweeney, Kate Connor, Klaus Dinkel, Michael Hamacher, Mark O. Cunningham, David C. Henshall, Tobias Engel

## Abstract

Resistance to anti-seizure medications (ASMs) remains a major clinical challenge in the management of epilepsy. Neuroinflammation has been implicated as a contributing mechanism and, consistent with this, antagonists of the ATP-gated P2X7 receptor (P2X7R) have been shown to enhance ASM efficacy in animal models. It remains unclear, however, if P2X7R antagonism is effective in human models of drug-resistant epilepsy. Here, we assessed the seizure-suppressive potential of the P2X7R antagonists AFC-5128 and JNJ-47965567 using electrophysiological recordings in acute resected brain slices from patients with epilepsy using artificial cerebrospinal fluid containing low Mg²⁺ and high K⁺ to evoked seizure-like events. P2X7R antagonists alone did not suppress seizure-like activity. When co-administered, however, P2X7R antagonists significantly enhanced the anti-seizure efficacy of carbamazepine and lorazepam. Notably, P2X7R antagonist treatment lowered Interleukin-1β levels, and betaine, a drug targeting Interleukin-1β release, mimicked the effects of P2X7R antagonists when combined with carbamazepine. Finally, mice with microglia-specific P2X7R depletion showed improved responsiveness to carbamazepine, suggesting P2X7R-mediated effects are partially mediated via their functions in microglia. These findings suggest that P2X7R-based therapies may represent an effective add-on approach for treating drug-resistant seizures associated with neuroinflammation.

## Introduction

Epilepsy, a heterogenous group of brain diseases characterized by the occurrence of spontaneous seizures, affects over 70 million people worldwide (Thijs et al., 2019). Common etiologies of epilepsy include genetic alterations and acquired brain insults such as traumatic brain injury, status epilepticus (SE), and brain tumors (Aronica et al., 2023; Dwivedi et al., 2024; Pitkanen et al., 2015).

First-line treatment in epilepsy is via anti-seizure medications (ASMs). These are effective, however, in only 70% of patients and have no proven disease-modifying potential. They can also cause serious side effects (*e.g*., fatigue, dizziness, headaches, memory impairment) (Klein et al., 2024; Loscher et al., 2022). Drug refractoriness remains one of the most significant challenges in epilepsy management (Kwan et al., 2011). Several pathological mechanisms have been implicated in the development of drug resistance, including alterations in neurotransmitter receptor expression or localization, blood-brain barrier (BBB) dysfunction, structural changes such as neurodegeneration, axonal sprouting, synaptic reorganization, gliosis, and neuroinflammation (Bazhanova et al., 2021; Costagliola et al., 2022; Loscher et al., 2020; Perucca et al., 2023; Tang et al., 2017). Among these, neuroinflammation has emerged as a particularly compelling contributor to pharmacoresistance and epileptogenesis, with studies showing that inflammatory mediators such as interleukin-1β (IL-1β) and High Mobility Group Box Protein 1 (HMGB1) increase neuronal excitability, alter glutamatergic and GABAergic neurotransmission, and impair responsiveness to ASMs (Vezzani et al., 2019).

Increasing evidence suggests that the ionotropic purinergic P2X7 receptor (P2X7R) plays a key role in the development and maintenance of neuroinflammation that promotes hyperexcitable neuronal networks in the brain (Beamer et al., 2021). The P2X7R is activated by high concentrations (µM - mM) of extracellular ATP (eATP) that occur when tissue is damaged. Upon activation, P2X7Rs form a non-selective cationic channel permeable to Na⁺, K⁺, and Ca²⁺. These receptors are widely expressed across the brain but are particularly abundant in immune cells such as microglia, where they contribute to the release of pro-inflammatory mediators including IL-1β (Alves et al., 2024; Illes et al., 2017; Kaczmarek-Hajek et al., 2018). P2X7Rs have been implicated in multiple pathological processes relevant to epilepsy, including aberrant synaptic plasticity and neurogenesis, neurodegeneration, and neuroinflammation (Andrejew et al., 2020; Klein et al., 2018; Sperlagh et al., 2014). In preclinical epilepsy models, antagonism of P2X7Rs has been shown to modulate seizure severity during SE and to suppress spontaneous seizures in epilepsy (Amhaoul et al., 2016; Amorim et al., 2017; Engel et al., 2012; Jimenez-Pacheco et al., 2016; Jimenez-Pacheco et al., 2013; Mamad et al., 2023; Rozmer et al., 2017). Notably, recent work in mouse models has demonstrated that P2X7Rs contribute to unresponsiveness to several ASMs, and that P2X7R antagonism can overcome inflammation-induced drug refractoriness (Beamer et al., 2022).

It remains unclear whether these beneficial effects will translate to humans. Recent studies indicated that P2X7R antagonism may attenuate epileptiform activity in human induced pluripotent stem cell (iPSC)-derived cells (Kesavan et al., 2026). Resective surgery for epilepsy not only provides an effective treatment option for patients with drug-resistant seizures but also offers a unique opportunity to directly test the efficacy of novel therapeutics in ex vivo brain tissue from pharmacoresistant individuals that retains disease-associated cellular and network alterations (Jones et al., 2016; Morris et al., 2021). Although increased P2X7R expression has been reported in resected brain tissue from patients with temporal lobe epilepsy (TLE) (Jimenez-Pacheco et al., 2016; Jimenez-Pacheco et al., 2013), the impact of P2X7R antagonism on epileptiform activity in human epileptic tissue has not yet been investigated.

Here, we show that combined treatment with the ASM carbamazepine or anticonvulsant lorazepam, P2X7R antagonists effectively reduce epileptiform-like activity in resected brain tissue from epilepsy patients. This effect was recapitulated by betaine, a drug that inhibits the release of IL-1β (Bhatt et al., 2024; Zhang et al., 2023). We further demonstrate that microglia-specific P2X7R deletion enhances responsiveness to carbamazepine during SE in mice, suggesting that effects are, at least in part, mediated via P2X7Rs expressed on microglia.

Together, these findings provide translational evidence from patient tissue that targeting P2X7R-mediated inflammatory pathways may represent a promising adjunctive therapeutic strategy for pharmacoresistant epilepsy.

## Materials and methods

### Patient tissue analysis

Selected patients suffered from drug-resistant epilepsy or brain tumors with and without seizures. For tumor patients, tissue was obtained from the peritumoral area (see **Supplementary Table 1** for patient demographics).

For electrophysiological studies, resected tissue was immediately submerged into cold carbogenated (95% O_2_ and 5% CO_2_) transport solution containing (in mM): 205 sucrose, 10 glucose, 3 KCl, 2 MgSO_4_, 2 CaCl_2_ 2 H_2_O, 24 NaHCO_3_, 1.25 CaCl_2_•2H_2_O) and transported to the laboratory. The tissue was then dissected into 400 µm thick slices on a vibratome (Zeiss Hyrax V50) in carbogenated transport solution. Slices were then transferred to interface chambers continuously bubbled and perfused with artificial cerebrospinal fluid (ACSF; in mM): 126 NaCl, 3 KCl, 1.25 NaH_2_PO_4_, 1 MgSO_4_, 1.2 CaCl_2_ 2H_2_O, 10 glucose, 24 NaHCO_3_), and allowed to recover for 1 h before start of recordings. Field recordings were obtained using borosilicate glass capillaries pulled with a micropipette puller (Sutter Instrument Co. Model P-97 Flaming) filled with ACSF with a resistance of 3-10 MΩ positioned in grey matter of the slice. The glass electrodes were connected to an NL100AK headstage, and the signals were amplified using an NL104 pre-amplifier (Digitimer, Hertfordshire, UK), band-pass filtered at 0.5 Hz to 500 Hz using NL125 Filter module (Digitimer, Hertfordshire, UK), and digitized using a CED Micro 1401 (Cambridge Electronic Design Ltd., Cambridge, UK) interfaced with a Windows 10 PC. The data were sampled at 5 kHz and processed using Spike2 (version 10.02, Cambridge Electronic Design Ltd., Cambridge, UK). Seizure-like events (SLEs) were defined as event lasting longer than 10 s, characterized by superimposed high-frequency, low amplitude activity followed by low-frequency rhythmic discharges resembling the clonic phase of a seizure. The SLEs were induced by perfusing slices with modified ACSF (mACSF) containing 8 mM K^+^ and 0.2 mM MgSO_4_ (high K^+^ low MgSO_4_). Pharmacologic experiments began 40 min after SLEs had appeared to allow for stabilization (baseline). Parameters analysed: event rate (SLEs / min); peak amplitude [peak (mV) considering the smooth field potential shift]; event duration [duration (s), measured from start up to two-thirds recoveries of the slow field potential]; and the power spectrum of the last 10 min of each treatment (from 0.5 Hz to 100 Hz). Values from individual events were averaged within each analysis epoch for every parameter. Drug effects were then quantified by normalizing the parameter values obtained during each drug-treatment epoch to their respective baseline values. Wash-out at the end of recordings confirmed that tissue remained excitable.

### Drug treatments in human tissue

Drug concentrations were taken from the literature and used at the following concentrations: This included the P2X7R antagonists AFC-5128 (i.e., 100 nM, 200 nM and 300 nM) and JNJ-47965567 (100 nM) (Kesavan et al., 2026) both made in DMSO and diluted in ACSF. The final concentration of DMSO was 0.0003% for AFC-5128 and 0.0001% JNJ-47965567. The IL-1β release blocker Betaine (1 nM) (Bhatt et al., 2024; Zhang et al., 2023) was made (Zhou et al., 2009) in ddH2O and then diluted in ACSF and the ASM carbamazepine (CBZ) 50 µM (Jandova et al., 2006) and lorazepam (100 µM) were made in polietilenglicol 400 and diluted in ACSF.

### Animals and mouse model of status epilepticus (SE)

All animals were housed in a controlled biomedical facility on a 12-hour light/dark cycle at 22 ± 1°C and humidity of 40-60% with food and water provided ad libitum. Studies to determine the effects of P2X7R deletion on microglia were carried out in 8-10 weeks old male and female mice using the tamoxifen-inducible Cre line B6.129P2(Cg)-Cx3cr1tm2.1(cre/ERT2)Litt/WganJ (*Cx3cr*) (JAX stock #021160) crossed to mice where the murine exon 2 of the *P2rx7* gene is flanked with loxP sites *(P2rx7^fl/fl^*) generated by the European Conditional Mouse Mutagenesis (EUCOMM) Program [P2rx7tm1a(EUCOMM)Wtsi] to obtain conditional mice with a *P2rx7* deletion in microglia (*P2rx7^-/-^*-M) (Alves et al., 2024). Tamoxifen treatment was applied for a period of 5 consecutive days by an i.p. injection of tamoxifen once daily (40 mg/kg; prepared in 10% of 100% ethanol and 90% of peanut oil; volume injection = 100 µl). Animals negative for Cre but homozygous for loxP were used as controls (*P2rx7*^fl/fl^). All mice received the same tamoxifen treatment regime.

SE was induced as described previously (Alves et al., 2024). Once fully anesthetized (5% induction, 1-2% maintenance), mice were placed in a stereotaxic frame and a midline scalp incision was performed to expose the skull. A guide cannula (coordinates from Bregma: AP = −0.94 mm, L = −2.85 mm) and three electrodes (Bilaney Consultants, Sevenoaks, UK), one on top of each hippocampus and with the reference on top of the frontal cortex, were fixed in place with dental cement. An Xltek recording system (Optima Medical, Guildford, UK) was used to record electroencephalogram (EEG). Following a recovery period of approximately 1 h post-surgery, SE was induced via a microinjection of 0.3 µg kainic acid (KA) in 0.2 µl Phosphate-buffered saline (PBS) (Sigma-Aldrich, Dublin, Ireland) into the right basolateral amygdala into awake, hand-restrained mice. To analyze seizure severity, EEG data were uploaded onto Labchart7 software (AD Instruments) as before (Beamer et al., 2022). EEG total power (µV^2^) is a function of EEG amplitude over time and was analyzed by integrating frequency bands from 0 - 50 Hz. Power spectral density heat maps were generated within LabChart7 (spectralview), with the frequency domain filtered from 0 - 40 Hz and the amplitude domain filtered from 0 - 50 mV.

### Elisa

The murine interleukin-1β (IL-1β) ELISA kit (Bio-Techne Ireland) was used according to the manufacturer’s instructions, where a wash comprises 3x washes of wash buffer/well (0.05% Tween-20 in PBS). Briefly, a clear 96-well plate was coated with capture antibody (50 µl per well) before incubating overnight at RT. The plate was then washed before blocking with reagent diluent (RD: 1% BSA in PBS) (150 µl per well, 1 h, RT). Standards were prepared and serially diluted (0-1000 pg/ml). Following washing, standards and samples were applied (50 µl per well) before further incubation (1 h, 37 °C). The plate was then washed again before incubating with detection antibody (50 µl per well, 1 h, 37 °C) Further washing followed this incubation before the addition of streptavidin-HRP (1:40 in RD, 50 µl per well, 20 min, RT). The plate was washed again before incubating with tetramethylbenzidine (1:1 H202, 50 µl per well, 20 min, RT, protected from light). Stop solution (H2SO4, 50 µl per well) was then applied to terminate the reaction before reading the absorbance of standards and samples at wavelengths of 450 nm and 540 nm using a plate reader. Il-1β concentration was interpolated from the standard curve, then normalized to milligrams of total protein concentration in tissue. Data is presented as n-fold of control samples.

### Statistical analysis

Statistical analysis of data was carried out using GraphPad Prism 8 and STATVIEW software (SAS Institute, Cary, NC, U.S.A). Data are presented as means ± standard error of the mean (SEM). One-way ANOVA parametric statistics with *post hoc* Fisher’s protected least significant difference test was used to determine statistical differences between three or more groups. Unpaired Student’s t-test (parametric) or non-parametric Mann-Whitney was used for two-group comparison. Normality and lognormality test were used to verify the normal distribution between groups. Significance was accepted at \**p* < 0.05.

## Results

### P2X7R antagonism potentiates the effects of anti-seizure medications (ASMs) in reducing epileptiform-like activity in resected patient tissue

Studies to date have demonstrated that P2X7R antagonism influences seizure generation and severity in preclinical models or human iPSC models (Engel et al., 2021; Fischer et al., 2016; Jimenez-Pacheco et al., 2016; Jimenez-Pacheco et al., 2013; Kesavan et al., 2026). To determine whether P2X7R antagonism also modulates epileptiform activity in a more translational model, we performed electrophysiological recordings from acute brain slices prepared from resected tissue obtained during epilepsy surgery. This included patients with TLE, Frontal lobe epilepsy (FLE) as well as patients with brain tumors (with and without seizure activity) (see **Supplementary Table 1**). Immediately after surgical removal, tissue was immersed in oxygenated high sucrose ACSF and processed for slice electrophysiology, followed by incubation in ACSF to assess treatment responsiveness under ex vivo conditions (**Fig. 1A**).

**Figure 1.**
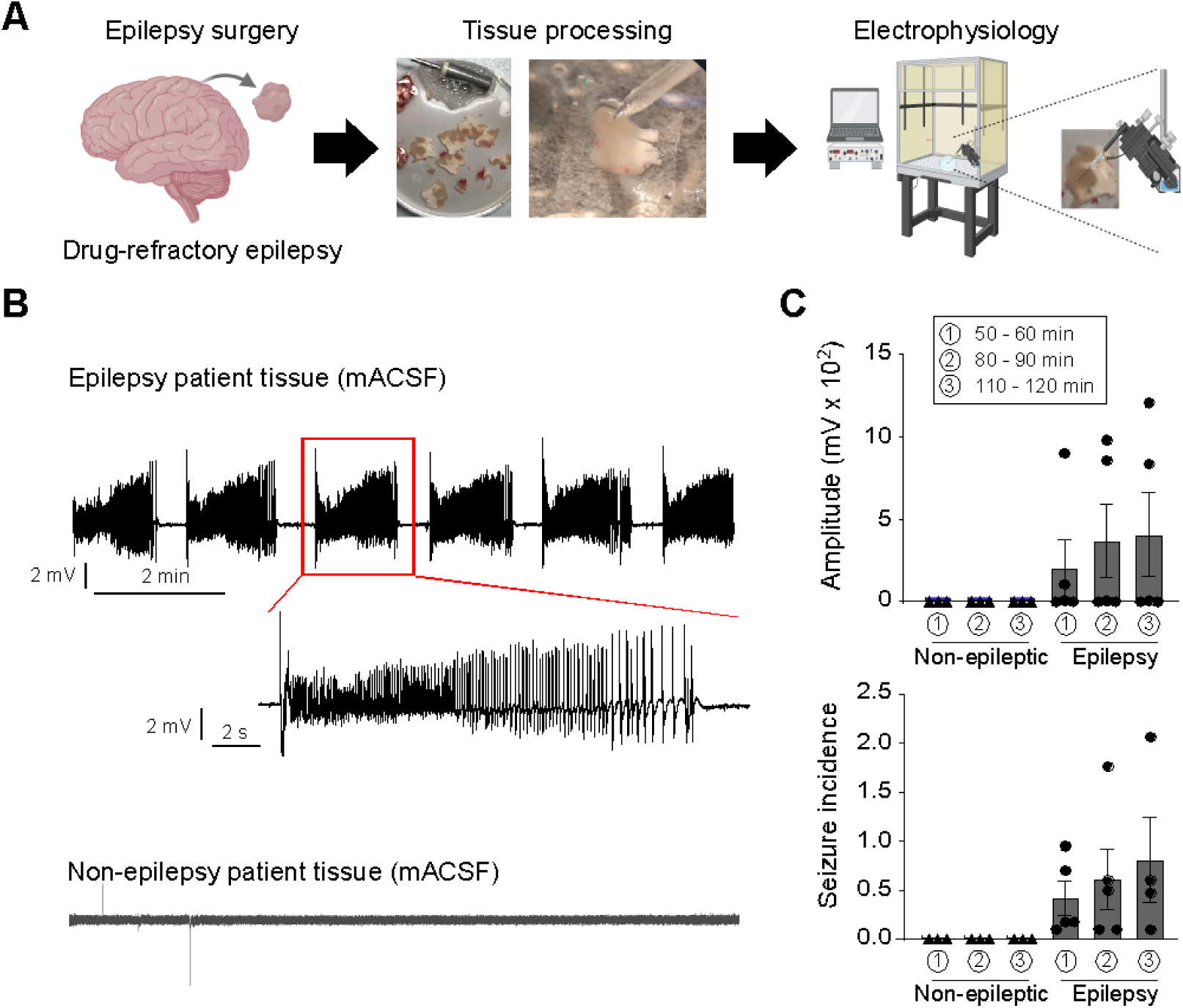
*Human epilepsy model. **A.** Schematic of the experimental design.* **B.** Representative EEG recordings from resected brain slices obtained from patients with and without epilepsy. Treatment with low Mg²⁺ and high K⁺ induced epileptiform activity only in brain slices from patients with epilepsy. **C.** Quantification of seizure incidence and signal amplitude in resected brain slices from patients with and without epilepsy treated with low Mg²⁺ and high K⁺ (N = 3 patients without epilepsy and N = 6 patients with epilepsy).

Seizure-like activity was induced using modified ACSF (mACSF) containing high K⁺ and low Mg²⁺ (**Fig. 1B**). Typically, seizure-like activity began approximately 40 min after incubation of tissue samples in mACSF, with seizure-like events (SLEs) exhibiting typical high-amplitude, high-frequency spiking usually lasting approximately 30 seconds (**Fig. 1B,C**). Brain slices treated with mACSF also presented increased power spectrum, amplitude and SLE frequency during the time, while SLE length did not show any significant change over time. Seizure activity normally persisting at least for a 120-minute recording period (**Supplementary Fig. 1A-C**). Notably, SLEs were only present in tissue sections obtained from patients who had previously experienced epileptic seizures, including patients with TLE and those in whom tumors were the underlying pathology. No SLEs were observed under our experimental settings in patients with tumors but without epilepsy (**Fig. 1B,C**).

Next, we sought to determine whether P2X7R antagonists on their own reduce evoked seizures in resected brain tissue. To address this, brain sections were first incubated with mACSF. Once a stable seizure baseline had been established, the P2X7R antagonist AFC-5128 was applied to the slices at different concentrations (100 nM, 200 nM, and 300 nM). Seizures were recorded for 40 min, followed by a 40 min washout period (**Supplementary Fig. 2A**). Suggesting no effect on seizure activity, treatment with AFC-5128 resulted in a slight increase in amplitude, power spectral density, and SLE incidence, consistent with the gradual increase in seizure severity observed over time in the resected tissue. In contrast, SLE duration was slightly reduced (**Supplementary Fig. 2B,C**).

Previous research has shown that P2X7Rs contribute to drug refractoriness during seizures and that P2X7R antagonism can overcome unresponsiveness to ASMs (Beamer et al., 2022; Engel et al., 2012; Fischer et al., 2016; Kesavan et al., 2026). To test whether P2X7R antagonism potentiates the effects of ASMs in our human brain slice model, brain slices were first incubated with the ASM carbamazepine, which inhibits voltage-gated sodium channels and is frequently used in clinical practice (Maan et al., 2026). This was followed by combined incubation with carbamazepine and the P2X7R antagonist AFC-5128 (Beamer et al., 2022) (**Fig. 2A**). Notably, while carbamazepine alone produced only mild seizure suppression at the dose used (in terms of SLE incidence, duration, and amplitude) compared with baseline, the combined application of carbamazepine and AFC-5128 reduced both the number and duration of SLEs (**Fig. 2B,C**).

**Figure 2.**
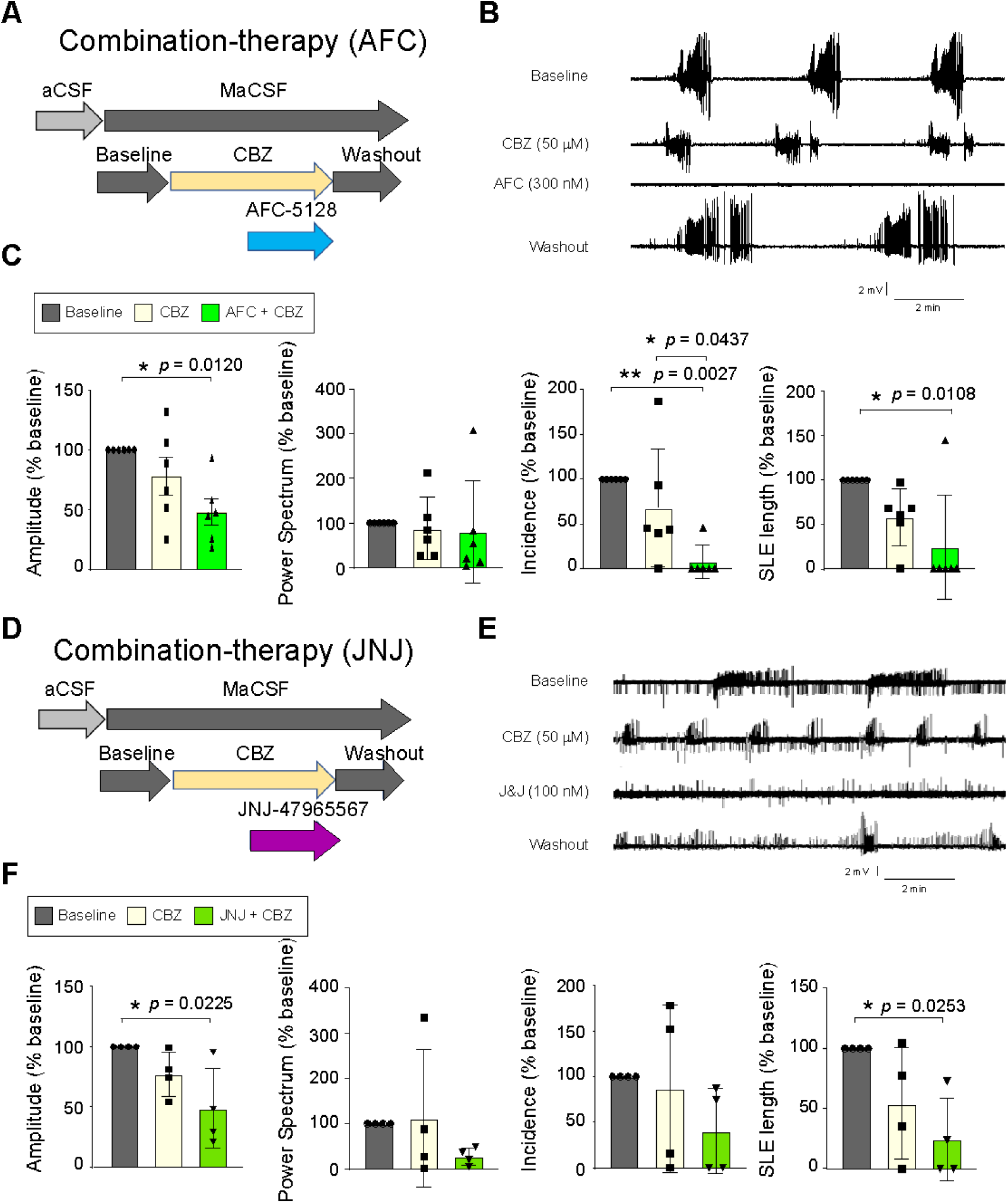
*Effect of P2X7R antagonism on epileptiform activity in resected brain slices.* **A.** Experimental design. Brain slices were incubated with mACSF for 30 min, followed by carbamazepine for 40 min and subsequently AFC-5128 (300 nM) for 40 min. **B.** Representative EEG recordings from brain slices following treatment with carbamazepine and AFC-5128. **C.** EEG amplitude, power spectrum, SLE incidence, and SLE duration during treatment with AFC-5128 (300 nM; N = 6). **D.** Experimental design. Brain slices were incubated with mACSF for 30 min, followed by carbamazepine for 40 min and subsequently JNJ-47965567 (100 nM) for 40 min. **E.** Representative EEG recordings from brain slices following treatment with carbamazepine and JNJ-47965567. **F.** EEG amplitude, power spectrum, SLE incidence, and SLE duration during treatment with JNJ-47965567 (100 nM; N = 4).

To confirm our results with AFC-5128, we tested a second allosteric P2X7R antagonist, JNJ-47965567 (Bhattacharya et al., 2013; Jimenez-Pacheco et al., 2016) (**Fig. 2D**). Consistent with our previous experiments, while carbamazepine alone produced only weak effects on seizure activity, the combined application of carbamazepine and JNJ-47965567 significantly reduced SLE duration, and amplitude compared with baseline (**Fig. 2E,F**).

Next, to determine whether the effects of P2X7R antagonism are specific to carbamazepine or whether seizure-suppressive effects are also observed with ASMs with distinct mechanisms of action, brain slices were treated with lorazepam, a benzodiazepine commonly used for the acute management of seizures that enhances GABAergic signalling through positive allosteric modulation of the GABA_A_ receptor (Goldschen-Ohm, 2022) (**Supplementary Fig. 3A**). Of note, mice overexpressing P2X7Rs are refractory to lorazepam (Beamer et al., 2022) and P2X7R antagonists have been shown to potentiate effects of lorazepam during drug-refractory SE (Engel et al., 2012). Similar to the results observed with carbamazepine, treatment with lorazepam alone at the selected dose (100 µM) had no effect on seizure suppression. In contrast, combined treatment with lorazepam and the P2X7R antagonist AFC-5128 reduced SLE incidence, duration, and amplitude (**Supplementary Fig. 3B**), indicating that the seizure-suppressive effects of P2X7R antagonism are not limited to a single ASM but may have broader applicability across distinct anti-seizure mechanisms.

Taken together, although P2X7R antagonists alone had no obvious seizure suppressive effects in our human slice model, P2X7R antagonism potentiated the effects of the ASM carbamazepine and lorazepam when applied in combination, suggesting that P2X7R antagonists may serve as an effective add-on therapy for drug-refractory seizures and epilepsy.

### P2X7R antagonism reduces IL-1β release in resected patient tissue, while betaine mimics the seizure-suppressive effects of P2X7R antagonism

Data from our group recently showed that genetic deletion of P2X7R from microglia reduces IL-1β levels in the hippocampus of mice following SE (Alves et al., 2024). Next, we investigated the effects of P2X7R antagonists of IL1 levels in the human model. Stimulation of slices with mACSF resulted in an increase in IL-1β in patient tissue while the P2X7R antagonist AFC-5128 reduced IL-1β levels to control levels (**Fig. 3A**). Notably, and further supporting a role for IL-1β in contributing to drug-refractoriness, treatment with betaine, an inhibitor of IL1β release, mimicked the effects of P2X7R antagonists on seizures in resected patient tissue, reducing carbamazepine-unresponsive evoked seizures (**Fig. 3B-D**). These findings suggest that IL-1β may be generated by epileptiform activity in this human brain model and its attenuation by P2X7R antagonists may contribute to the anti-seizure mechanism.

**Figure 3.**
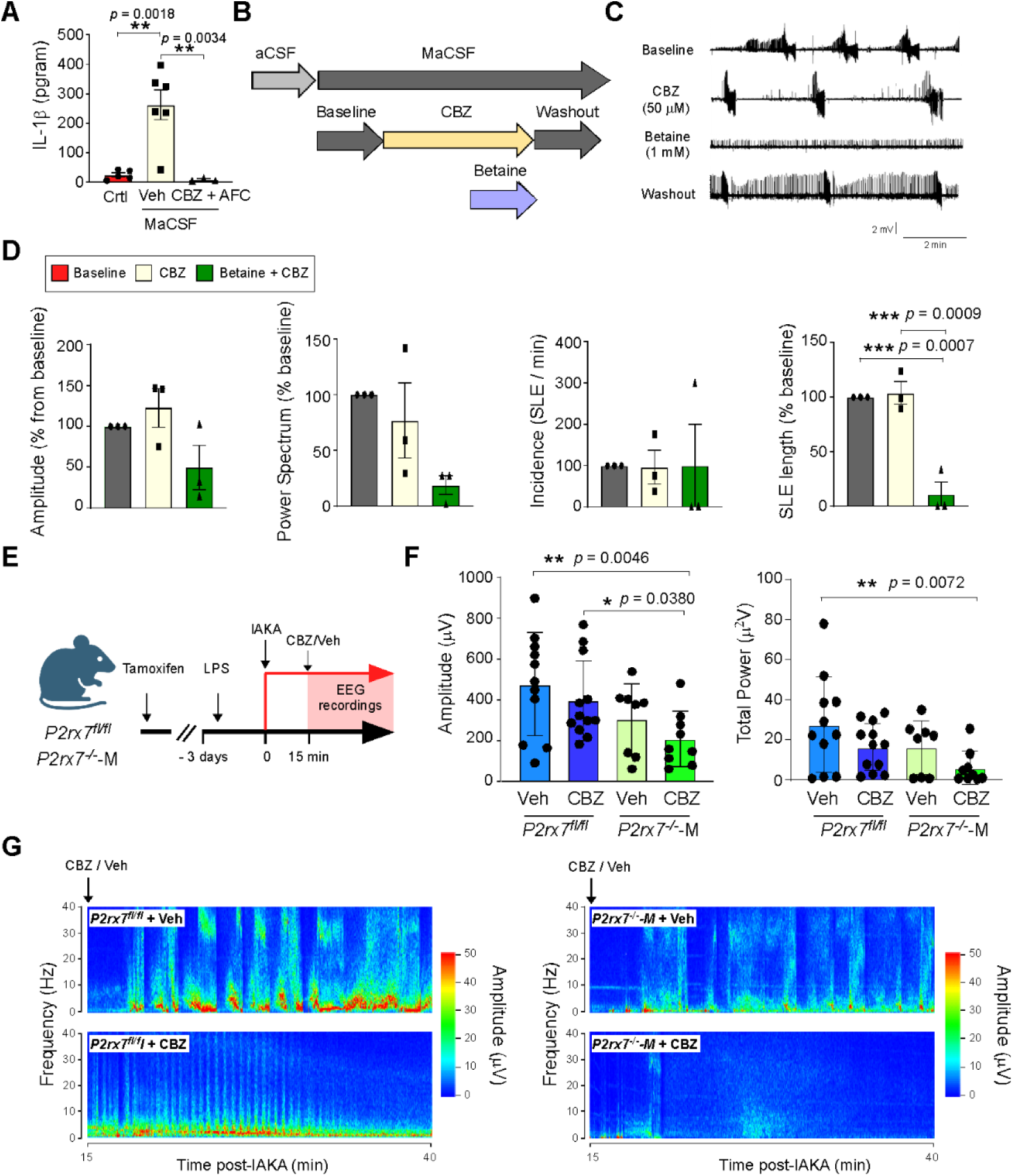
*Effects of IL-1β modulation on epileptiform activity in resected brain slices.* **A.** IL-1β concentrations in resected brain slices under control conditions, following treatment with mACSF, or following treatment with carbamazepine and AFC-5128. **B.** Experimental design. Brain slices were incubated with mACSF for 40 min, followed by carbamazepine for 40 min and subsequently Betaine for 40 min. **C.** Representative EEG recordings from brain slices following treatment with carbamazepine and Betaine. **D.** EEG amplitude, power spectrum, SLE incidence, and SLE duration during treatment with Betaine (N = 3). **E.** Experimental design. Three days following LPS treatment, *P2rx7*^fl/fl^ and *P2rx7*^-/-^-M mice were subjected to IAKA. Mice were then treated with carbamazepine or vehicle 15 min later, and cortical EEG was recorded for 25 min. **F.** Total EEG power and amplitude (N = 9 - 12 per group) from EEG recordings performed from the time of carbamazepine or vehicle treatment (15 min post-IAKA) until 40 min post-IAKA. **G.** Representative heatmaps for all four treatment groups from 15 to 40 min post-IAKA.

### Deletion of microglial P2X7Rs enhances the anticonvulsive efficacy of carbamazepine during seizures

P2X7Rs are highly expressed in microglia, including in epilepsy (Morgan et al., 2020), where they are thought to contribute to seizure generation and drug refractoriness via driving pro-inflammatory signalling (Beamer et al., 2022; Engel et al., 2021). Notably, mice with a microglia-specific deletion of P2X7Rs display reduced seizure severity during SE and develop a milder epilepsy phenotype (Alves et al., 2025; Alves et al., 2024).

To test whether mice with microglia-specific P2X7R knockout (KO) respond better to the ASM carbamazepine, tamoxifen-treated P2X7R KO microglial mice (*P2rx7⁻/⁻*-M) and corresponding controls (*P2rx7^fl/f^l*), were pre-treated with LPS (Beamer et al., 2022) to increase brain inflammation and subjected to IAKA-induced SE. Mice where then treated with carbamazepine or vehicle 15 min post-IAKA, time-point when mice experienced at least 5 min of seizure activity (**Fig. 3E**). EEG analysis revealed that the anti-convulsive effects of carbamazepine were more pronounced in microglial P2X7R KO mice (**Fig. 3F,G**), suggesting that microglial P2X7Rs contribute to unresponsiveness to carbamazepine.

## Discussion

Here, we report that P2X7R antagonism potentiates the anticonvulsant effects of ASMs in resected tissue from epilepsy patients. We demonstrate that P2X7Rs regulate seizure-induced IL-1β release and that blocking IL-1β release mimics the seizure-suppressive effects of P2X7R antagonists. Finally, we show using genetic techniques in mice that these effects are at least partly mediated by P2X7Rs located on microglia. Together, our results suggest P2X7R antagonists may be a suitable add-on therapy for drug-refractory epilepsy.

The main finding of our study is that P2X7R antagonism reduces seizures when administered in combination with ASMs in an epilepsy model using resected human tissue. While animal models remain the principal platform for identifying and testing new drug candidates for seizure control and epilepsy (Loscher et al., 2023), offering the advantage of enabling investigation of complex interactions from the molecular to the behavioral level, attrition rates following the preclinical phase remain high, even for compounds that initially show promise with the lack of efficacy being the main cause (Bialer et al., 2026). Resected tissue from patients, in turn, offers a unique opportunity to evaluate drug effects directly in human tissue, thereby providing a potential bridge between preclinical testing and clinical application (Nogueira et al., 2022). Testing in patient-derived tissue not only enables the evaluation of drug effects within a relevant disease context but also allows assessment of whether clinical candidates effectively target human proteins. This is particularly important for P2X7Rs, which exhibit marked species-specific differences in sequence and function (e.g., increased affinity of AFC-5128 to the human P2X7R (Bartlett et al., 2014; Fischer et al., 2016), underscoring the need for validation in human tissue.

The demonstration that P2X7R antagonism potentiated the effects of the ASMs carbamazepine and lorazepam is consistent with findings from animal studies. For example, P2X7R antagonism has been shown to potentiate the effects of lorazepam in the IAKA mouse model (Engel et al., 2012) and of the ASM carbamazepine in the maximal electroshock seizure test in mice (Fischer et al., 2016). Notably, P2X7R overexpression has been linked to drug refractoriness in mice during SE including both lorazepam and carbamazepine (Beamer et al., 2022). The present findings also complement a recent study using human iPSCs that showed that P2X7R antagonism via AFC-5128 and JNJ-47965567 reduced seizure-like activity in an inflammation-primed drug-resistant model (Kesavan et al., 2026).

P2X7R-mediated drug refractoriness appears to be linked to the receptor’s role in driving pro-inflammatory processes in the brain (Engel et al., 2021). This is consistent with our data showing that microglia-specific P2X7R KO mice exhibit an improved response to carbamazepine during SE. Previous studies have implicated neuroinflammation, including IL-1β signaling, in drug refractoriness (Roseti et al., 2015). Microglia respond rapidly to neuronal hyperactivity by adopting a pro-inflammatory phenotype and releasing cytokines that directly influence neuronal excitability, and pronounced microglial activation has been reported in resected tissue from patients with drug-refractory epilepsy (Choi et al., 2009). Although several ASMs can exert anti-inflammatory effects, ongoing microglial activation may limit their efficacy in drug-refractory epilepsy. In this context, targeting microglial P2X7Rs represents a complementary therapeutic strategy. Consistent with previous reports showing that P2X7R antagonism reduces IL-1β following SE (Alves et al., 2024; Engel et al., 2012) and in patient tissue (Cowley et al., 2026), our findings support the idea that inhibition of microglial P2X7Rs dampens seizure-induced neuroinflammation, thereby restoring the ability of ASMs to effectively engage their neuronal targets and suppress seizures.

The mechanism by which P2X7R-dependent IL-1β signaling impairs ASM efficacy remains to be fully established. IL-1β can modulate neuronal ion channels, including voltage-gated sodium channels, thereby increasing excitability and reducing responsiveness to ASMs (Roseti et al., 2015). We therefore propose that seizure-induced activation of microglial P2X7Rs amplifies pro-inflammatory signaling, including IL-1β release, which interferes with ASM target engagement. Conversely, pharmacological inhibition or genetic deletion of microglial P2X7Rs reduces neuroinflammation, restoring ASM efficacy and enhancing seizure suppression. While our data suggest that effects on seizures are mediated, at least in part, through IL-1β, other P2X7R-dependent inflammatory mediators, including TNF-α and IL-18, may also contribute and warrant further investigation.

While our mouse findings identify microglia as the principal cell type mediating the pro-inflammatory effects of P2X7Rs, consistent with their high levels of P2X7R expression (Kaczmarek-Hajek et al., 2018), contributions from other glial cell types cannot be excluded. For example, P2X7Rs have also been shown to be expressed by astrocytes and oligodendrocytes (Zhao et al., 2021), both of which have been implicated in epilepsy and neuroinflammation (Onat et al., 2025; Vezzani et al., 2022). Future studies using cell type-specific deletion of P2X7Rs in these populations will be important to define their relative contributions to seizure generation, neuroinflammation, and drug refractoriness.

The present study explored the effects of P2X7R antagonism in combination with carbamazepine and lorazepam, two ASMs with distinct mechanisms of action. While carbamazepine acts primarily by blocking voltage-gated sodium channels, lorazepam acts as a positive allosteric modulator of the GABA_A_ receptor (Goldschen-Ohm, 2022; Maan et al., 2026). It remains unknown if the potentiating effects of P2X7R antagonism will extend to other ASMs.

The finding that, when applied alone, P2X7R antagonism has no obvious effect on epileptiform activity suggest that P2X7R targeting specifically on microglia may be more effective and that P2X7R antagonism is not inherently anticonvulsant, as previously shown (Fischer et al., 2016), but rather enhances seizure suppression when combined with ASMs (Thakku Sivakumar et al., 2024). These results further suggest that P2X7R antagonism may be most effective in the presence of an inflammatory background.

While IL-1β inhibition reproduced the seizure-suppressive effects of P2X7R antagonism, P2X7R remains the more attractive therapeutic target because it acts upstream of multiple inflammatory pathways and P2X7R antagonists have shown favorable safety and tolerability in clinical trials for other indications (Palmer et al., 2026; Timmers et al., 2018). By enhancing ASM efficacy, P2X7R antagonists may also allow lower ASM doses, potentially reducing treatment-associated adverse side effects.

There are a number of limitations to consider in the present study. While useful in demonstrating the translatability of novel treatments, resected tissue has its own limitations including disrupted neuronal networks, changes in the tissue due to surgical manipulation and slice preparation, lack of a BBB and possibly increased ATP release, main endogenous agonist of P2X7Rs. Nevertheless, demonstrating therapeutic efficacy directly in human epileptic tissue represents an important step toward clinical translation.

In conclusion, our findings identify P2X7R inhibition as a promising adjunctive strategy to overcome drug refractoriness in epilepsy. Importantly, demonstrating these effects directly in resected human epileptic tissue provides strong translational support and may help accelerate the clinical development of P2X7R-targeted therapies.

## Author Contributions

Conceptualization, A.F.M., B.G. and T.E.; methodology, A.F.M., B.G., M.M., J.K., A.L., MJ.S., A.M.OC. and T.E.; formal analysis, A.F.M., B.G., M.M., and T.E.; resources, V.H., N.D., A.B., F.M.B., M.A.F., J.C., D.F.OB., K.J.S., K.C., K.D., M.H., M.O.C., D.C.H. and T.E.; writing - original draft preparation, T.E.; writing - review and editing, All.; supervision, T.E.; project administration, T.E.; funding acquisition, J.K., D.C.H. and T.E. All authors have read and agreed to the published version of the manuscript.

## Ethics declaration

All animal experiments were performed in accordance with the principles of the European Communities Council Directive (2010/63/EU). Procedures were reviewed and approved by the Research Ethics Committee of the Royal College of Surgeons in Ireland (RCSI) (REC 1322) and Health Products Regulatory Authority (HPRA) (AE19127/P075). All subjects gave their informed consent for inclusion before they participated in the study. The study was conducted in accordance with the Declaration of Helsinki, and the protocol was approved by the Ethics Committee of Beaumont Hospital, Dublin (05/18).

## Disclosure

AI tools were used for language editing, grammar correction, and stylistic refinement of the manuscript. The authors utilized ChatGPT for these purposes and take full responsibility for the accuracy, content, and any changes made, ensuring all information aligns with research standards. This use is disclosed as per journal guidelines.

