## Supplemental Table 1 for "P2X7 receptor antagonism potentiates seizure-suppressive effects of anti-seizure medications in human resected epileptic brain tissue"

**Supplementary Table 1**: *Demographic and clinical characteristics of patients with epilepsy*

| **Experiment** | **Sex** | **Age** | **Seizure onset** | **Confirmed Seizures** | **Diagnosis** | **ASMs at time of surgery** | **Tissue specimen** |
| --- | --- | --- | --- | --- | --- | --- | --- |
| AFC | F | 52 | 13 | Yes | Bilateral TLE, HS Type 1 | BRIV, LTG | Lateral neocortex |
| mACSF | M | 46 | 19 | Yes | TLE | LEV, LCS, CEN | Lateral neocortex |
| AFC | M | 57 | 21 | Yes | FLE | CLOB, LTG, LEV, MID | Frontal cortex |
| mACSF | M | 47 | 37 | Yes | TLE | CLOB, ESLI, LTG | Lateral neocortex |
| AFC | M | 28 | 8 | Yes | TLE, bottom of sulcus FCD Type 2A | LTG, LAC, LEV | Lateral neocortex |
| mACSF | M | 22 | 17 | Yes | TLE | LAC, MID, VAL, ZON | Temporal neocortex |
| AFC | M | 37 | 37 | Yes | Grade 4 astrocytoma, isolated seizure | None | Peritumoural |
| AFC | F | 77 | 77 | Yes | GBM, collapse/query seizure | None | Temporal peritumoural cortex |
| AFC | F | 39 | 23 | Yes | TLE | BRIV, CLB, LCS, LTG | Lateral neocortex |
| mACSF | M | 50 | 36 | Yes | TLE | VPA, ZNS | Lateral neocortex |
| mACSF | F | 24 | 3 | Yes | FLE | ESLI, ZNS | Lateral neocortex |
| CBZ+  AFC | F | 39 | 28 | Yes | TLE, MTS+ (right side); epilepsy tissue secondary to stroke | CEN, ESLI, LSM | Lateral neocortex |
| CBZ + AFC | F | 30 | 21 | Yes | "MRI negative, PET positive" right TLE | ESLI, LCS | Lateral neocortex |
| CBZ + AFC | F | 40 | 23 | Yes | Right TLE | LTG, LCS | Lateral neocortex |
| LZ+AFC | M | 48 | 48 | yes | Grade 2 oligodendroglioma.  Acute symptomatic seizures. | LEV | Frontotemporal peritumoural cortex |
| CBZ + AFC | M | 69 | 69 | yes | Grade 3 oligodendroglioma.  Acute symptomatic seizures. | LEV | Parietal peritumoural cortex |
| LZ+AFC | M | 37 | 0.5 | Yes | Long standing TLE | BRV, CLB, LTG, LZP, MID, OXC, PB, VPA, ZNS | Left posterior quadrant TPO disconnection with IOM |
| CBZ+ AFC | F | 55 | 21 | Yes | Long-standing drug refractory TLE | BRV, CEN, LCM | Lateral neocortex |
| LZ+AFC; CBZ + Betaine | M | 35 | 19 | Yes | Long-standing drug refractory localisation related epilepsy, cause unknown | BRV, CEN, LTG, PER | Posterior left insula resection |
| CBZ + JnJ | M | 71 | - | Yes | Query localisation related epilepsy cause unknown | CEN, LTG | Right sided anterior temporal lobectomy |
| CBZ + JnJ; CBZ + Betaine | M | 36 | - | Yes | Medication refractory TLE | LEV, LCM, PHT | Lateral neocortex |
| mACSF | M | 38 | N/A | No | Non epilepsy/no seizure history | None | Lateral neocortex |
| CBZ + JnJ; CBZ + Betaine | M | 51 | 51 | Yes | Acute symptomatic isolated generalised tonic clonic seizure | LEV | Lateral neocortex |
| mACSF | F | 34 | N/A | No | Left frontal diffusely infiltrating grade 2 astrocytoma | - | Peritumoral cortex |
| CBZ + JnJ; LZ + AFC | M | 36 | 36 | Yes | Query isolated symptomatic seizure | LEV | Peritumoral cortex |
| CBZ + AFC | M | 48 | 48 | Yes | Glioblastoma | LEV, BD | Peritumoral cortex |
| mACSF | M | 54 | N/A | No | Glioblastoma | LEV, BD | Peritumoral cortex |

**Abbreviations:** ***BRIV***, **Brivaracetam**; ***CEN***, **Cenobamate; *CLB***, **Clobazam; *CLOB***, **Clobazam; *ESLI***, **Eslicarbazepine; *FCD***, **Focal Cortical Dysplasia; *FLE***, **Frontal Lobe Epilepsy; *GBM***, **Glioblastoma; *HGG***, **High-Grade Glioma;** *HS*, Hippocampal sclerosis; *IOM*, Intraoperative Monitoring; ***LAC***, **Lacosamide;** ***LCS***, **Lacosamide**; ***LEV***, **Levetiracetam; *LSM***, **Lacosamide; *LTG***, **Lamotrigine; *LZP***, **Lorazepam;** ***MID***, **Midazolam; *MTS***, **Mesial Temporal Sclerosis; *MRI***, **Magnetic Resonance Imaging; *OXC***, **Oxcarbazepine; *PB***, **Phenobarbital; *PER***, **Perampanel; *PET***, **Positron Emission Tomography; *PHT***, **Phenytoin;** *TLE*, Temporal lobe epilepsy; *TPO*, Temporo-Parieto-Occipital; ***VAL***, **Valproate; *ZNS***, **Zonisamide; *ZON***, **Zonisamide**
