## Supplemental Material for "P2X7 receptor antagonism potentiates seizure-suppressive effects of anti-seizure medications in human resected epileptic brain tissue"

**Supplementary Material**

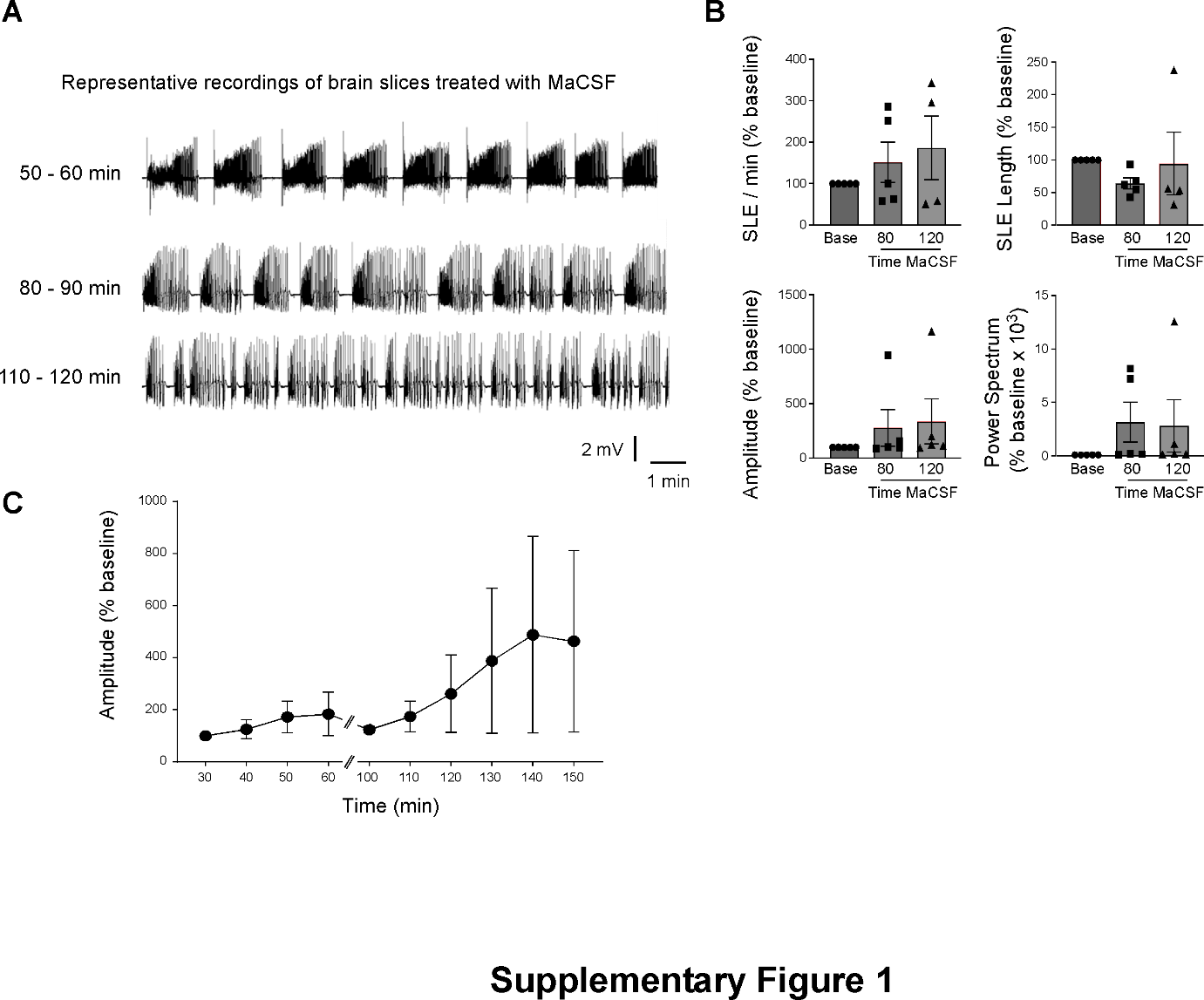

**Supplementary Figure 1.** *Effects of mACSF treatment on seizure-like events in resected patient tissue maintained in ACSF over time.* **A.** Representative EEG recordings from brain slices at different time points during incubation with mACSF. **B.** Quantification of SLE frequency, SLE duration, EEG amplitude, and power spectrum at different time points during incubation with mACSF, expressed as a percentage of baseline (20 - 30 min; N = 5). No significant changes over time were observed for any of the parameters analysed. **C.** EEG amplitude over 150 min in resected tissue from patients with epilepsy during incubation with mACSF (N = 5).

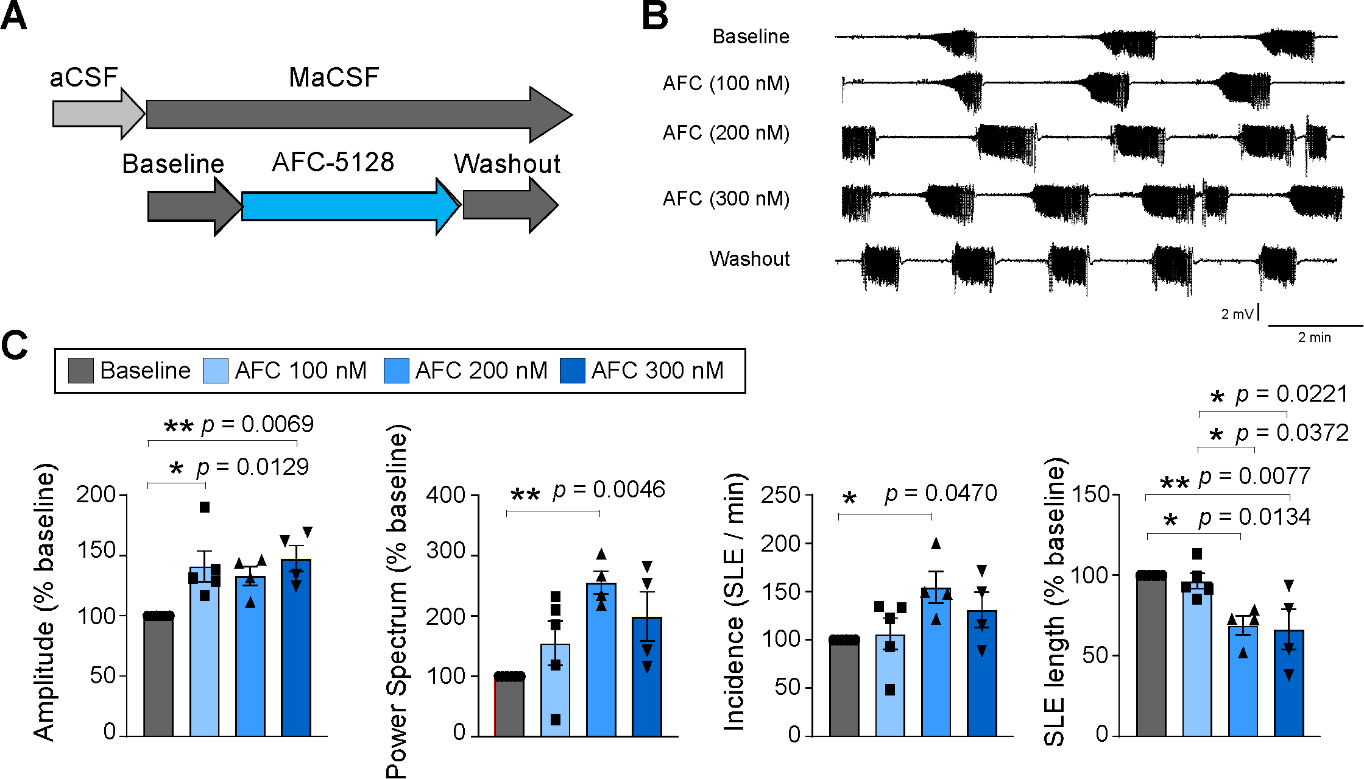

**Supplementary Figure 2.** *Effects of AFC-5128 on evoked seizures in resected tissue.* **A.** Experimental design. Brain slices were incubated with mACSF for 40 min, followed by incubation with AFC-5128 (100, 200, or 300 nM) for 30 min. **B.** Representative EEG recordings from brain slices following treatment with mACSF, AFC-5128 (100, 200, or 300 nM), and during drug washout. **C.** EEG amplitude, power spectrum, SLE incidence, and SLE duration during treatment with AFC-5128 (N = 5 for 100 nM AFC-5128 and N = 4 for 200 and 300 nM AFC-5128).

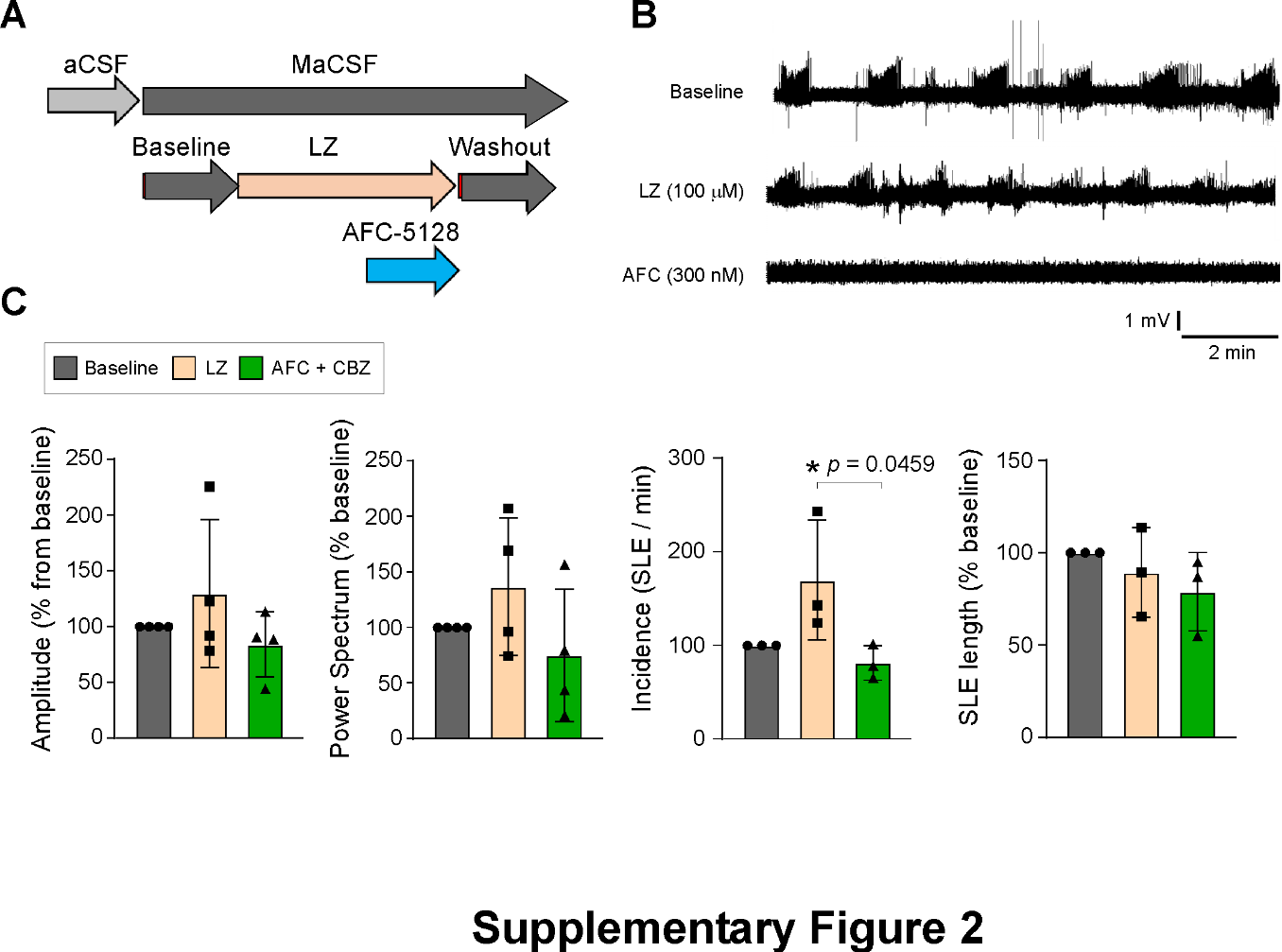

**Supplementary Figure 3.** *Effects of P2X7R antagonism on lorazepam-mediated modulation of epileptiform activity in resected brain slices.* **A.** Experimental design. Brain slices were incubated with mACSF for 40 min, followed by lorazepam alone (100 nM) for 30 min and subsequently by co-application of lorazepam (100 nM) and AFC-5128 (300 nM) for 40 min, followed by a 30 min washout period. **B.** Representative EEG recordings from brain slices following treatment with lorazepam and AFC-5128. **C.** SLE incidence, SLE duration, EEG amplitude, and power spectrum during treatment with AFC-5128 (N = 4).
